# B cells and HSV-specific antibodies respond to HSV-2 reactivation in skin

**DOI:** 10.1101/2020.07.08.192542

**Authors:** Emily S. Ford, Anton M. Sholukh, RuthMabel Boytz, Savanna S. Carmack, Alexis Klock, Khamsone Phasouk, Jason Shao, Raabya Rossenkhan, Paul T. Edlefsen, Tao Peng, Christine Johnston, Anna Wald, Jia Zhu, Lawrence Corey

**Affiliations:** Vaccine and Infectious Diseases Division, Fred Hutch Cancer Research Center, Seattle WA; Division of Allergy and Infectious Diseases, Department of Medicine, University of Washington, Seattle WA; Department of Laboratory Medicine and Pathology, University of Washington, Seattle WA; Department of Epidemiology, University of Washington, Seattle WA

## Abstract

Tissue-based T cells increasingly have been shown to be important effectors in the control and prevention of mucosal viral infections – less is known about tissue-based B cells. We demonstrate that B cells and antibody-secreting cells (ASCs) are present in skin biopsies of persons with symptomatic HSV-2 reactivation. CD20^+^ B cells are observed in inflammatory infiltrates at greatest density at the time of symptomatic reactivation; HSV-2-specific antibodies to HSV-2 surface antigens are also detected. The concentrations of HSV-2-specific antibodies in tissue biopsies vary over the course of HSV-2 reactivation and healing, unlike serum where concentrations remain static over time. B cells and HSV-specific antibody were rarely present in biopsies of unaffected skin. Investigation of serial biopsies over the course of lesion healing suggests that B cells follow a more migratory than resident pattern of infiltration in HSV-affected genital skin, in contrast to T cells. Together, these observations may suggest a functional and distinct role of tissue-based B cells in the local immune response to HSV-2.

## Introduction

In immunocompetent hosts, after initial exposure at the site of non-intact epithelialized skin or mucosa, herpes simplex virus 2 (HSV-2) infection establishes chronic latent infection of the associated dorsal root ganglion (1, 2). The spectrum of clinical disease among infected persons is broad; some experience asymptomatic seroconversion while others have 10 or more recurrences a year. While the mechanisms underlying viral reactivation and symptomatic disease are not known, the spectrum of clinical manifestations at the site of reactivation is thought to depend primarily on the host adaptive immune response, rather than intrinsic viral factors (3, 4). As suggested by human-based studies and supported by mathematical modeling and animal studies, cell-mediated immunity, including tissue-resident memory CD8^+^ and infiltrating or resident memory CD4^+^ T cells are important factors in local viral control (5–10).

The role of humoral immunity in the human response to HSV-2 reactivation is less clear. Robust titers of binding and neutralizing antibodies are commonly detected in the blood, but are not associated with the frequency of HSV-2 symptoms in natural infection (11, 12) or after vaccination (13–15). Vaccine-induced antibody to gD was not protective against primary infection in HSV-2 seronegative persons (16). One proposed theory is that a dominant antigen, gD, leads to production of antibodies that are binding and neutralizing, but not protective (17). Other proposed theories include that certain HSV-specific antibodies may function protectively in tissue near the site of viral release but circulating antibody levels do not reflect this tissue-based event (18), or that cell-to-cell spread of virus is sufficient for propagation and development of symptomatic reactivation even in the presence of effective neutralizing antibodies, but the concentration of such antibodies is below the threshold of efficacy (19).

Animal models indicate that B cells are important to protection from HSV-2 and other viral infections. B cell-deficient mice rapidly develop severe disease after HSV-2 genital challenge (6, 20). B cells are present in cutaneous inflammatory infiltrates induced by vaccinia virus in mice, potentially due to lymphocyte recruitment by VCAM-1 (21). In HSV-2-exposed mice, B cells infiltrate into the vaginal mucosa in response to immunization with latency-deficient HSV-2. These B cells are associated with intraluminal vaginal secretion of HSV-specific IgG2 and IgA during viral challenge, but not at other times (22). Jiang et al (2017) showed HSV-specific antibody and ASCs within the trigeminal ganglia of HSV-1-infected mice. These observations, in concert with the description of memory lymphoid clusters in mice infected with HSV-2, raised the possibility that B cells and ASCs may function outside of their previously understood role only in lymph tissue. Wilson et al. (2019) define an IgM^+^ population of B cells that migrate from the peritoneum and accumulate in inflamed skin in mice in response to chronic inflammation (23). In humans, the role of B cells during mucosal HSV-2 infection has not been well-studied. From single cell dissociation studies, B cells have been identified in normal skin, but are very rare (24, 25). With histologic confirmation of their presence in the tissue extrinsic to blood vessels, antibody-secreting cells (ASCs) have been identified in human foreskin (26). B cells have been observed, albeit infrequently, in inflammatory infiltrates related to HSV (27), and their presence in the skin has been confirmed by single-cell and RNAseq technology (25, 28, 29). Whether B cells and ASCs are actively involved during the immune response to HSV-2-reactivation in humans in non-mucosal skin, therefore, is unknown.

We utilized our repository of sequential genital skin biopsies from persons with HSV-2 to explore whether B cells and ASCs are present in genital skin biopsies taken during HSV-2 reactivation and clinical quiescence. HSV-2 specific antibody is present in tissue and the concentration varies over time suggesting the possibility of local antibody production. In a time of rapidly-expanding information from single-cell dissociation studies, this histologic information confirms the importance of spatially-informative methods, and urges further investigation into the role of B cells and ASCs in local inflammatory processes.

## Results

Of 16 participants, 14 were women; the median age was 50 years (range 18-66) (**Supplementary Table 1)**. All 16 were seropositive for HSV-2 and 6 also had antibody to HSV-1. The most common site of biopsies was the buttock (N=10) (**Supplementary Table 1**). Biopsies were performed during a clinically evident HSV-2 lesion and during healing, for a total of 5 or 6 biopsies per person. A control site biopsy of uninvolved skin was performed at the time of the lesion or 8 weeks after healing from either the arm (N=7) or the contralateral genital skin (N=9). There were no complications associated with serial biopsies (5, 8).

### Histology of B cells in tissue by immunofluorescence

B cells were identified by dual immunofluorescent staining of CD20 and CD79b in 47 of 65 punch biopsies of active and early healing HSV-2 lesions from 11 of the 16 persons. Per person, a median of 5 of 6 biopsies were seen to have B cells (range 2-6 of 6); in most persons B cells were present in all of the biopsies from the HSV-2 lesion site. CD20 and 79b were chosen for IF due to their relative abundance on the B cell surface (CD20 is expressed at 6-fold higher density than CD19 (30)) and consistent performance in this assay. Dual staining was employed to assist in identification of non-specific antibody binding. The number of CD20/CD79b^+^ cells present in each biopsy section varied by person and timepoint. In general, B cells were more common during the lesion and early healing, similarly to T-cell infiltration (5). They were infrequent in biopsies during clinical quiescence (present in 3 of 11 biopsies at 8 weeks) and were detected in only 3 of 16 control site biopsies either from uninvolved genital skin or the arm (**Figure 1A**).

**Figure 1.**
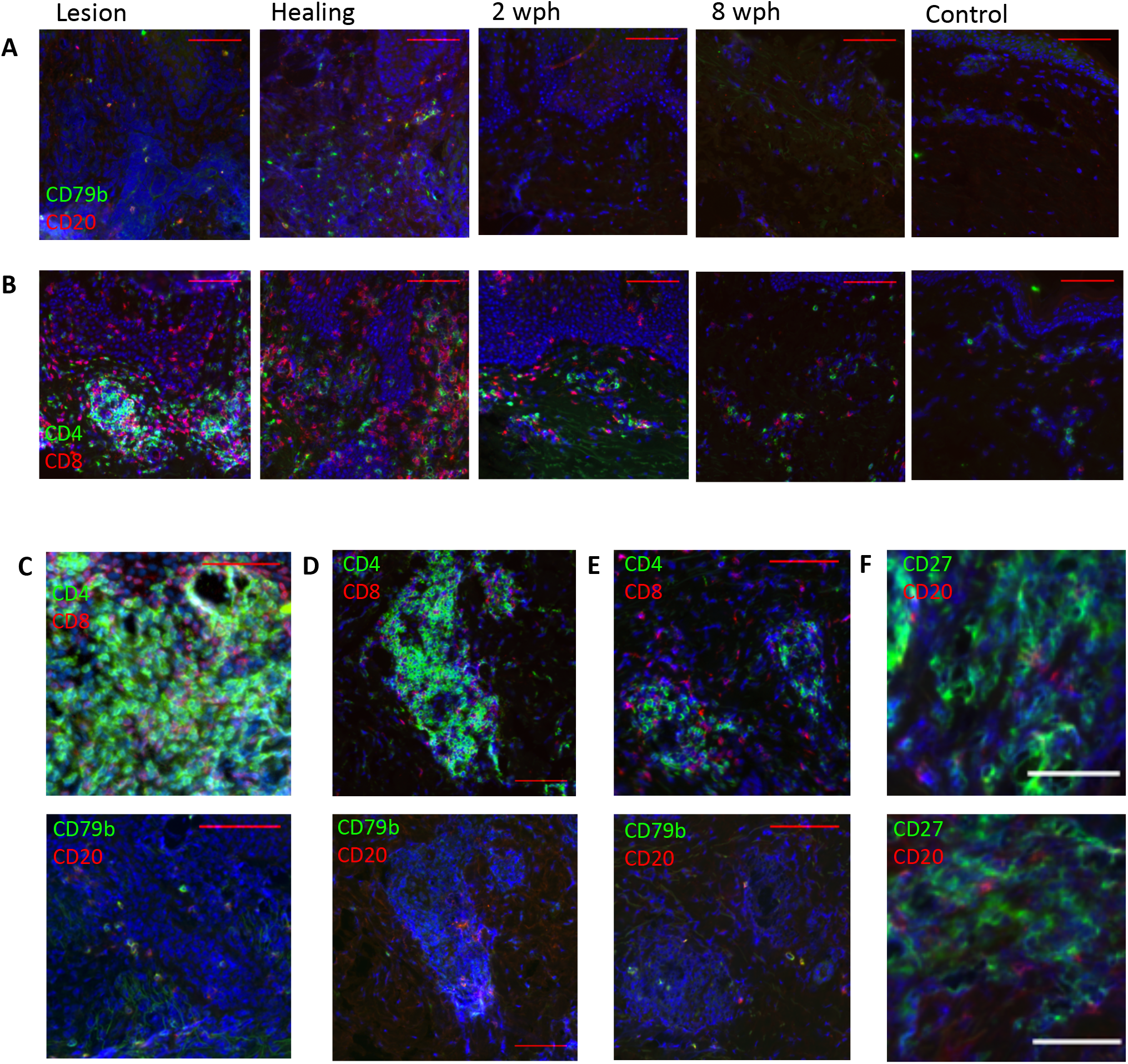
CD20^+^CD79b^+^ B cells infiltrate genital skin during the tissue-based immune response to HSV-2 reactivation. (**A**) B cells CD20 (red) and CD79b (green) (bottom) in serial genital skin biopsies during and after HSV-2 lesion from a single subject. Scale bars represent 100 μm. There were no B cells detected in this individual in the 8 wph or control biopsies. (**B**) T cells: CD4 (red) and CD8 (green) (top) in serial genital skin biopsies during and after HSV-2 reactivation from a single subject. (**C-E**) Histologic findings of T and B cells by IF (same antibodies as **A**,**B**) of skin biopsies during HSV-2 reactivation and healing. (**C**) Diffuse intradermal infiltration by T cells (top) corresponds with scattered infiltration by B cells (bottom). (**D**) B cells (bottom) are present in a dense cluster of T cells (top). (**E**) B cells present with T cells in intra-dermal glandular structures. (**F**) CD27 (green) and CD20 (red) from a lesion biopsy with no evidence of co-staining. Brightness of all images is increased by 30% for visualization.

Histologically, B cells were identified by CD20 IF in the upper dermis (within 500 μm of the surface epithelium) and were never found within the dermal-epidermal junction (DEJ) (**Figure 1A**), where CD8^+^ T cells are often found (**Figure 1B**) (5), or the multiple layers of the epidermis, consistent with histology in mice (23). CD20/CD79b^+^ cells were most often observed to be distributed within areas of immune cell infiltration (**Figure 1C**), but were also identified similarly located in dense groups of T cells by staining of serial sections (**Figure 1D**), suggesting in vivo interactions between the two subpopulations of cells. T cells are also often observed to be present near hair follicles or skin-localized glandular structures and this was likewise occasionally observed of B cells (**Figure 1E**). CD27, a canonical marker of memory B cells, is more strongly expressed by activated T cells and thus is abundant in skin biopsies during HSV-2 reactivation and healing. Co-staining of this marker may therefore be unreliable in biopsies with robust T-cell infiltration, but in B-cell containing biopsies from 5 individuals little to no overlap in staining of CD20 and CD27 was observed in B cells, suggesting that the majority of observed CD20^+^ cells are not memory B cells (**Figure 1F**).

To investigate the presence and distribution of antibody-secreting cells (ASCs), fluorescent in situ hybridization (FISH) directed at IgG mRNA (ACD Bio) was performed in 57 biopsies from 10 persons to identify cells with dense expression of immunoglobulin transcripts. Both class-switched B cells and ASCs contain IgG mRNA, however, ASCs are known to express up to 1000-fold more immunoglobulin mRNA per cell than B cells (31, 32). High intensity, cytoplasmic punctate fluorescence indicative of ASCs was observed in 48 of the 57 tested biopsies (**Figure 2A, 2B**). Histologically, these IgG RNA^+^ cells followed tissue distribution patterns described above seen with CD20^+^ B cells. No IgG RNA^+^ cells were observed within the epithelial cell layers or in the DEJ in any biopsy. All IgG RNA^+^ cells observed were present in the dermis. No IgG RNA^+^ cells by FISH were identified in any of the control biopsies.

**Figure 2.**
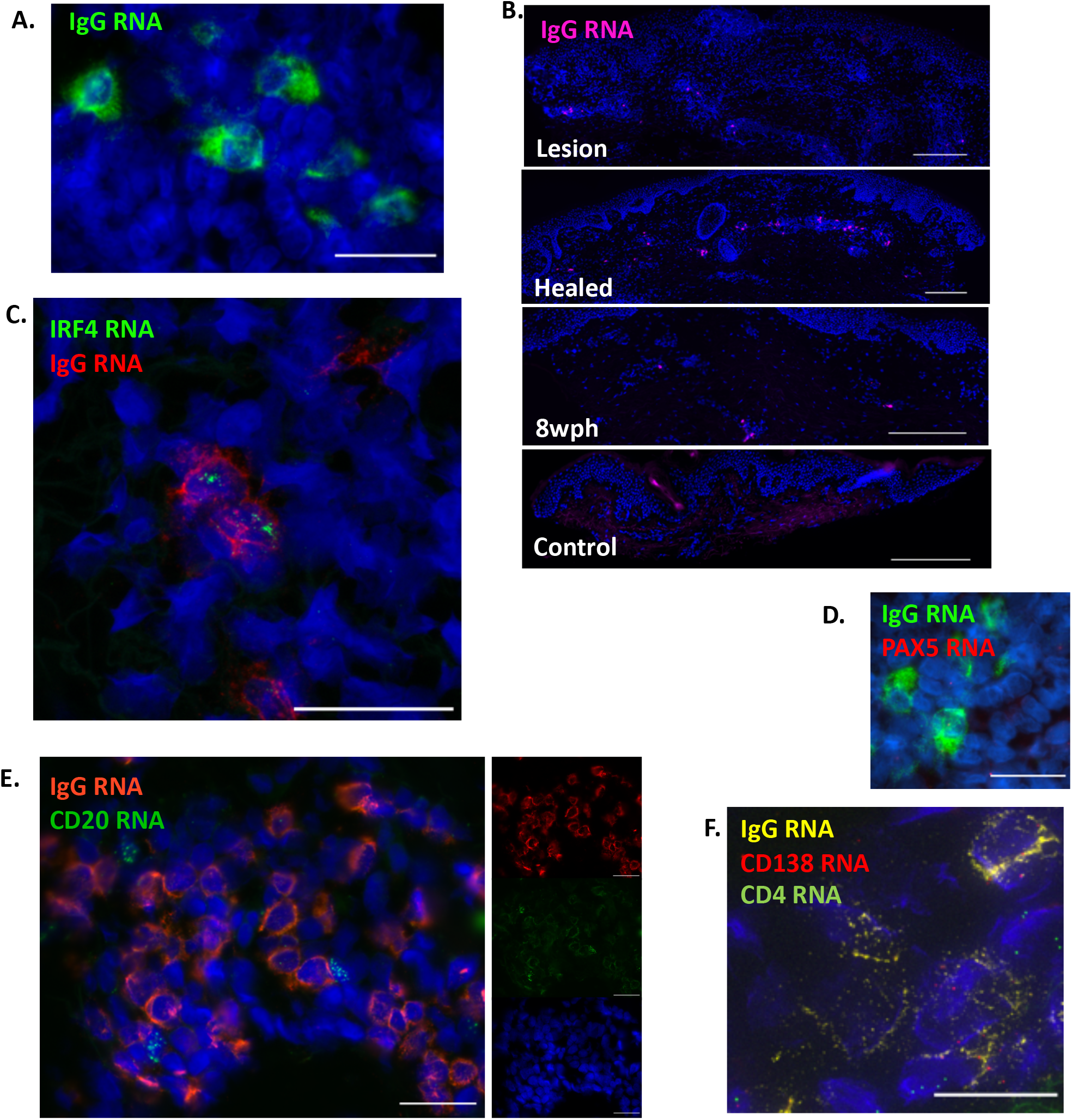
Cells expressing high level of IgG RNA infiltrate genital skin during the tissue-based immune response to HSV-2 reactivation. (**A**) IgG-producing cells by FISH. IgG mRNA (green) in a healing biopsy of HSV-2 reactivation. 40x magnification by oil immersion microscopy. Scale bar is 25μm. **(B)** IgG-producing cells by FISH over time showing distribution within the upper dermis and lymphocyte clusters. Images were obtained at 20x. Scalebar is 250μm. (**C**) IgG mRNA (red) and IRF4 (green) in a healing biopsy of an HSV-2 lesion. Note two cells with lower amplitude of IgG that are IRF4-. 60x magnification by confocal microscopy. Scale bar is 25 μm. (**D**) PAX5 (red) and IgG (green) co-expression in one of five IgG RNA^+^ cells in a small cluster. 40x magnification by oil immersion fluorescent microscopy, scale bar is 25 μm. **(E)** IgG (red) and CD20 (green) adjacent cells without co-expression. 40x magnification by oil immersion fluorescent microscopy, scale bar is 25μm. **(F).** IgG (yellow) and CD138 (red) co-expression by deconvolution microscopy (60x). CD4 (green) is seen on adjacent cells. Scale bar is 10μm. Brightness in A and D is increased by 30% for visualization and consistency.

Further confirmation of the presence of ASCs was pursued by dual FISH with expression of transcriptional factor IRF4 which is upregulated in ASCs by 5–10-fold compared to naïve and memory B cells (33). Nuclear IRF4 production was frequently observed to be co-localized with IgG transcripts by confocal microscopy (**Figure 2C**). Taken together, these findings strongly indicate the presence of IgG-producing ASCs during HSV-2 reactivation and skin healing.

B-cell-specific activator protein (PAX5) is a transcription factor that is highly expressed in naïve and memory lineage B cells but is downregulated by 4-5 fold in ASCs (33). Therefore, colocalization of a high level of PAX5 with low level of IgG transcript can indicate memory B cells. In genital skin, we observed rare PAX5 transcripts occasionally in cells where IgG transcripts were detected at high abundance (**Figure 2D**), but this was uncommon. The overall frequency of PAX5 detection was low, and PAX5 nuclear signal was also seen in cells not identified to have IgG transcription (IE non-B cell lineage cells). Minimal PAX5 expression in cells containing a high level of IgG transcripts supports that these cells are indeed ASCs. To confirm this phenotype, co-expression of CD20 and CD138 was also studied. Transcripts of CD20 were visualized both with and without expression of IgG, though more often was seen independently (**Figure 2E**), and CD138 signal was observed in some, but not all cells expressing a high level of IgG transcripts. Due to the abundant presence of CD138 in skin, deconvolution microscopy was used to confirm that CD138 and IgG signal were concomitantly present in single cells (**Figure 2F**). This is consistent with the interpretation that these cells are antibody-secreting.

While there are no validated markers to indicate tissue residence or migration of B cells, one important consideration is whether these observed B cells and ASCs have migrated into the tissue or whether they are present in tissue samples due to localization in small capillaries. By use of IF co-staining with von-Willenbrand Factor (vWF), which is expressed on the surface of endothelial cells, while IgG RNA^+^ cells identified by FISH were often seen near small capillaries, when localized diffusely in the upper dermis they were not contained within vascular structures (**Figure 3A**). Small vessels containing IgG RNA^+^ cells were identified in the lower dermis in biopsies from multiple individuals and these accounted for some of the apparently clustered IgG RNA^+^ cells in these biopsies. Clusters of IgG RNA^+^ cells were also observed that did not have evidence of associated endothelial cells (**Figure 2D, Fig. 3B, 3C**). Some clusters were found to contain T cells, as described previously (7, 8, 34), and ASCs were often observed within these clusters of T cells. Clusters were located in the dermis and contained both CD8^+^ (**Figure 3B**, by FISH and IF over serial sections) and CD4^+^ (**Figure 3C,** by dual-probe FISH) T cells. Notably, these immune cell clusters have not been detected in any of the 27 control biopsies collected either from the arm or contralateral genital area. We also did not see any B cells or ASCs in vascular structures in control biopsies. Thus, dermal-infiltrating and capillary-associated B cells and ASCs may all be related to HSV-2 reactivation in genital skin.

**Figure 3.**
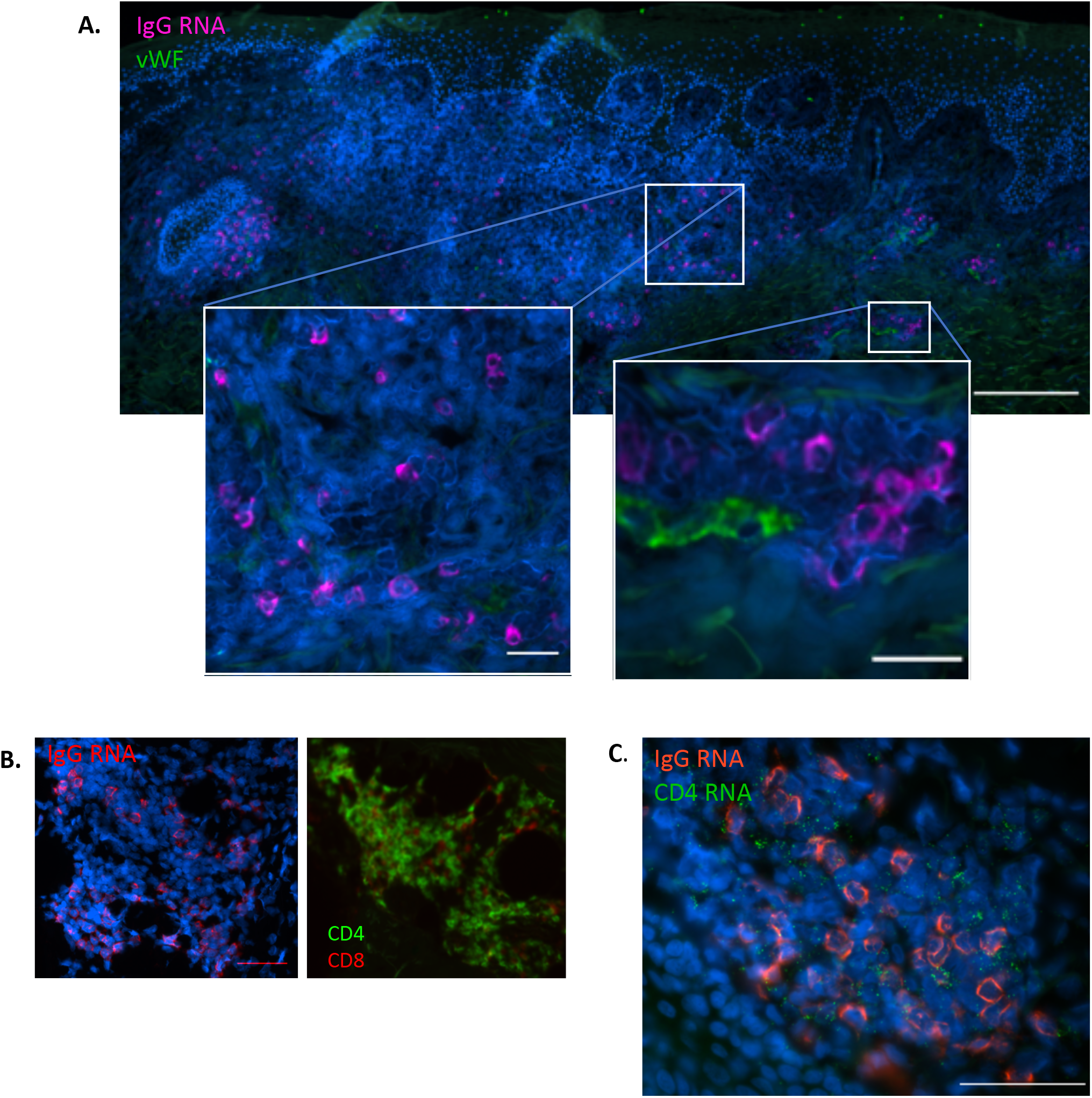
IgG RNA co-staining with other tissue structures and cell types. (**A**) Antibody-secreting cells are found infiltrating tissue without relation to small capillaries, adjacent to small capillaries, and within small capillaries. IgG+ cells by FISH (magenta) are found infiltrating in tissue (left) as well as proximate to and within (right) small capillaries identified by vWF IF (green), in a biopsy of healing HSV-2 reactivation. Scale bar in large image is 200μm, bars in insets are 25μm. (**B**) IgG-producing cells by FISH (left) clustered in area of dense T-cell infiltration (CD4 (green) and CD8 (red) (right), shown are serial sections of the same biopsy. Scale bar is 50μm. (**C**) Clustered IgG (red) and CD4 (green) by FISH. Scale bar is 50μm.

### Kinetics of B-cell response during HSV-2 reactivation and skin healing

Overall, the magnitude and density of the B-cell and ASC infiltrate (**Figures 4A and B**) were greatest during the acute phases of reactivation; i.e., during clinical lesion and early healing. HSV-2 was detected by PCR at the time of lesion biopsy in 12 of 14 patients in whom this was tested, and in the first healing biopsy in patients 7 and 16. In most persons, the highest density of CD20^+^ B cells was observed in biopsies when genital lesions were present; median density of 18.4 cells/mm^2^ (N = 8, range 2.6-46.0 cells/mm^2^). In three persons, the density of CD20^+^ cells peaked at the time of healing, about 10 days after lesion onset (**Figure 4A**), with a median of 16.1 cells/mm^2^ (range 1.5-37.4 cells/mm^2^). To determine whether the peak density of CD20+ B cells was higher at either active/early healing or later healing time points, the maximum of these biopsies for each person were compared. By Wilcoxon signed rank test, the maximum CD20^+^ cell density of ulcerative or healing lesions was higher than the maximum of the 2, 4, or 8 weeks post healing biopsies (p = 0.003). In three persons (1, 7 and 10), few B cells were seen in any biopsies.

**Figure 4.**
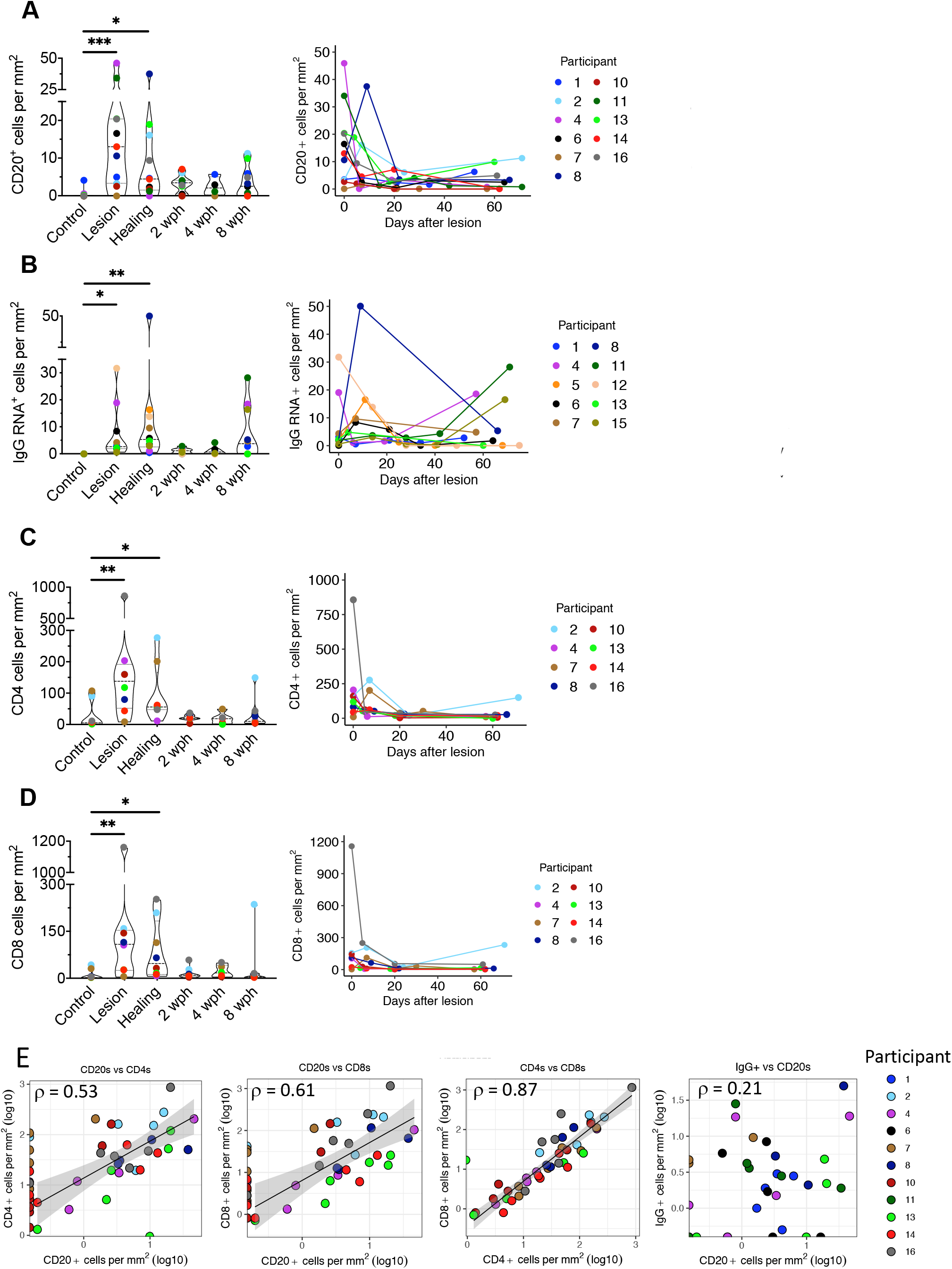
Migration kinetics of B and T cells into genital skin during HSV-2 reactivation and tissue healing by type of biopsy (left) and over time (right). Density of (**A**) CD20^+^ cells by IF, (**B**) IgG RNA^+^ cells by FISH, (**C**) CD4^+^ T cells by IF and (**D**) CD8^+^ T cells by IF in tissue over time and shown by persons with cell density peak at lesion (middle) or healing (right) time points. Asterisks indicate statistical testing by Friedman test with Dunn’s corrections for multiple comparisons. *P<0.05; **P<0.01, ***P<0.001. Each graph on the right presents cell counts from genital area biopsies over time, as measured from the identified symptomatic lesion. Lack of correlation between B and T cell subsets and between CD20^+^ and IgG RNA^+^ cells is shown in (**E**), Participants are labeled by color, multiple biopsies from genital and control areas from each participant are included in each graph.

In three persons (4, 11 and 15), we noted an increase in IgG RNA^+^ cells between the 4 and 8 weeks post-healing biopsies. Similarly, the CD20^+^ B cell density increased at 8 weeks post healing in patients 2 and 13. We speculated that this could be related to subclinical viral reactivation, albeit proof of this was not present in any participant by HSV PCR from tissue. In contrast to CD20+ B cells, by Wilcoxon signed rank test, the maximum IgG^+^ cell density of lesion or healing was no higher than the maximum of the 2, 4, or 8 weeks post healing biopsies (p = 0.2). In those three persons with a peak of IgG^+^ cells at 8 weeks post healing, the corresponding tissue antibody concentrations varied over time (4, 11, and 15); two had a peak at the post-healing time and one peaked during the lesion, none of these persons had a corresponding peak in antibody titer at the 8 week time point.

The density of CD20^+^ and IgG RNA^+^ cells was considerably lower than T-cell density at the site of HSV-2 reactivation (**Figure 4C and 4D**). Similarly to past reports (8, 27), CD4^+^ and CD8^+^ T cells were found at highest density during a symptomatic lesion in the majority of persons. As with CD20^+^ cells, by Wilcoxon signed rank test, the maximum CD4^+^ and CD8^+^ cell density of lesion or healing was higher than the maximum of the 2, 4, or 8 weeks post healing biopsies (p = 0.008, p = 0.04). We evaluated the relationship between B- and T-cell density in individual lesion biopsies (**Figure 4E**). Biopsies with the highest T-cell density tended to be those with higher B cell density. The Spearman coefficient for a correlation between CD20^+^ cells and CD4^+^ and CD8^+^ T cell subsets was 0.53 and 0.61, respectively, suggesting a potential relationship between these cell types, compared to 0.21 between CD20^+^ cells and IgG^+^ cells. For comparison, the Spearman coefficient between CD4^+^ and CD8^+^ T cells was 0.87 (**Figure 4E**). The simultaneous migration and spatial clustering of CD4^+^ and CD20^+^ cells in genital skin during HSV-2 reactivation and healing suggest that these lymphocyte subsets may be influenced by similar recruitment mechanisms.

### Detection and kinetics of HSV-2 specific antibodies in genital tissue

Antibody (IgG) was detectable by IF to human IgG in tissue. As with B cells, IgG was detected to the greatest extent in the upper dermis of lesion and early healing tissue biopsy sections and was less detectable in biopsies of uninvolved sites (**Figure 5A**). In a biopsy of an active HSV-2 vesicle, IgG was detectable co-localized with HSV antigen (**Figure 5B**). Fluorescence was not detected with an isotype control, isolated secondary antibody or if the tissue was blocked with 5% human serum prior to IF (**Supplementary Figure 1**). We therefore hypothesized that IgG detected in skin during HSV-2 reactivation (**Figure 5**) is produced by an antigen-specific process, such as by locally-migrated B cells, and would therefore be present at higher concentration than non-HSV-specific antibody.

**Figure 5.**
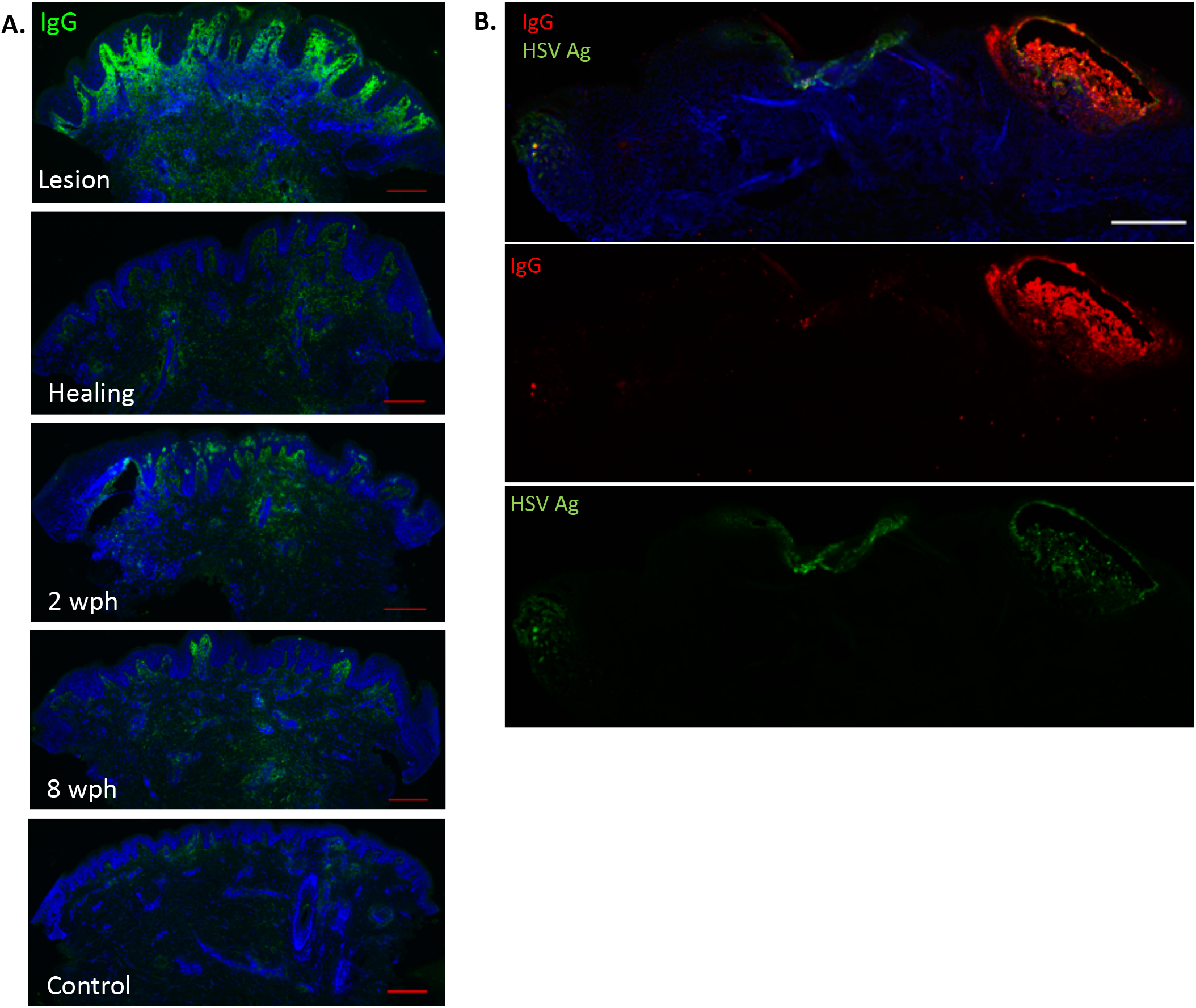
Antibody is detectable during the tissue-based immune response to HSV-2 reactivation and the amount is variable over time. (**A**) Immunofluorescent detection of IgG (green) in genital skin biopsies in one subject. Scale bar represents 250μm. Brightness is increased by 30% for visualization. (**B**) Spatial localization of in-tissue IgG vs HSV antigen by IF in biopsy of an active HSV-2 lesion (different subject). Antigen is seen in multiple locations with the highest density of IgG at the site of vesicle formation. Scale bar is 250μm.

We first investigated the trajectory of the systemic antibody response during HSV-2 reactivation and healing via sequential serum samples from two persons collected concurrently with skin biopsies (all other participants had a single blood draw at study entry, which may not have corresponded with their biopsy series). We used an antibody binding assay based on the Luminex platform to detect antibodies specific to the major HSV-2 surface glycoproteins gB2 and gD2. We included EBV gp350 and the stem of influenza hemagglutinin (Flu-HA) as highly prevalent, non-HSV targets. For all tested antigens, no variability in serum antibody level was detected over time as evidenced by overlapping titration curves and area under the curve (AUC) measurements (**Supplementary Figure 2** and **Supplementary Table 2**). This finding is consistent with earlier reports documenting the absence of systemic antibody response during local HSV-2 reactivation (35–38).

To investigate whether immunoglobulin observed in biopsies by IF was specific to HSV-2, we extracted immunoglobulin from sequential biopsies from 14 participants to measure IgG reactive to gB2, gD2, and total IgG. IgG reactive to gB2 and gD2 proteins was detectable in the majority of the lesion area biopsies but was not detected in many biopsies collected from the arm and contralateral skin (**Figure 6A, 6B**). When followed over time, the pattern of shifts in concentration of HSV-specific IgG (**Figure 6A, 6B**) and total IgG (**Figure 6C**) varied over time and by individual, similarly to B cells and ASCs. Over participants the amount of HSV-specific antibody varied by up to 30-fold. Within participants, the difference in the amount of HSV-specific antibody in different biopsies likewise varied substantially (gB2 median 3.4-fold, range 1.7-57.8, and gD2 median 6.1-fold, range 2.3-94.7). Some participants had a peak of HSV-2 specific IgG at the lesion, others had a peak during early healing, and the rest showed a peak of HSV-2-specific IgG at late healing. A higher concentration of total IgG was found at the lesion in biopsies from 6 participants and post-healing in 4 participants, the total amount of IgG in the control biopsies was generally low (**Figure 6C**). Conversely, there was no major change observed over the course of a lesion and healing in the levels of IgG specific to control antigens in skin extracts: EBV gp350 and Flu HA (**Figure 6D**) except participant 6 for whom IgG at the lesion was substantially higher than for subsequent time points. Antibodies to control antigens were observed in skin extracts of only a few persons.

**Figure 6.**
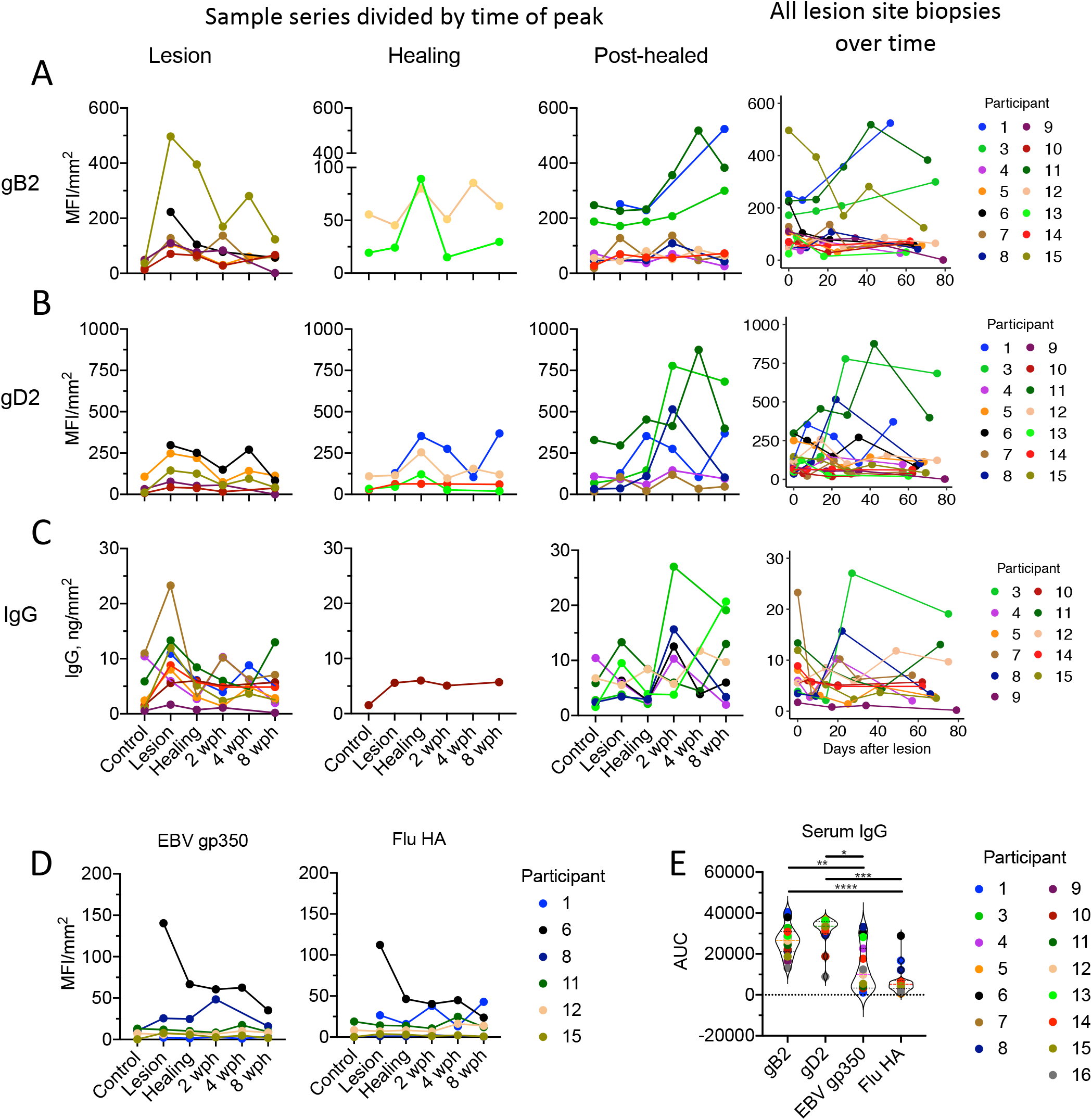
Levels of HSV-2 specific antibody and total IgG follow B-cell and ASC density. IgG levels are expressed as MFI units per mm^2^ of biopsy. Concentration (normalized to biopsy surface area) of (**A**) gB2, (**B**) gD2, (**C**) total IgG, and (**D**) control antigens EBV-gp350 and Flu-HA in tissue over time and shown by persons with antibody peak at lesion (1^st^ column) or healing (2^nd^ and 3^rd^ column) time points. If a person did not have a clear peak at one time point, they were included in multiple graphs. Levels of antibody over time (by number of days after lesion biopsy) is in the 4^th^ column. Missing data points (such as the arm control in person 6) correspond to biopsies with insufficient tissue to run the test. (**E**) The area under the curve in a serum measurement of antibody specific to each of these antigens from a single sample taken from each person at study entry.

When applied to serum samples obtained at study intake for all individuals, the amount of circulating antibody to gD2 (as measured by AUC) was similar for the majority of the participants while the amount of gB2-specific IgG was more varied among participants. Both gB2 and gD2-specific IgG were present at significantly higher levels than IgG against non-HSV antigens, although the difference in levels between HSV-2 antigens and EBV gp350 was less profound (**Figure 6E, Supplementary Figure 3 and Table 3**).

This discordance between HSV-specific and control antigens in skin and serum also supports in-tissue antibody production or an antigen-driven B-cell recruitment mechanism. While some of the discrepancy between control and HSV-specific antibody in tissue may be accounted for by its greater abundance in serum, that was not consistently true. For example, the level of EBV-gp350 IgG was similar to gB2 IgG in the serum of participant 13, whereas in skin EBV-gp350 IgG was not detected and gB2 IgG peaked at the healing time point (**Figure 6D**). In participant 8, EBV-gp350 IgG in serum was present at a higher level than gD2 and gB2-specific Abs, but in skin gD2 IgG peaked at 2wph while EBV gp350 IgG was increased at lesion time point. Interestingly, Flu-HA but not EBV IgG mirrored the dynamics of HSV-specific antibody in this participant. Together, these findings support multiple mechanisms of antibody infiltration into inflamed skin, including antigen-independent transudation from serum and antigen-dependent processes such as local production.

## Discussion

Here we have shown that B cells and ASCs are present in biopsy tissue from persons undergoing biopsies of active and healing skin lesions due to HSV-2 reactivation. Our studies clearly show B cells to be within the upper dermis and extrinsic to blood vessels in the weeks during and after HSV-2 reactivation. Spatially, we see two patterns of B-cell infiltration. In the first, ASC are present in the dermis at the site of HSV reactivation in variable density over time, and this coincides with the pattern of total immunoglobulin we see during genital lesions. The other pattern is the presence of CD20+ B cells in T-cell clusters. These B cells are mainly CD20+ cells without memory markers and were often not associated with Ig secretion by FISH staining; perhaps suggesting these are early stage B cells that might be participating in Ag processing or some as yet undefined function, such as regulation of inflammation. The lack of correlation between IgG+ and CD20 cells in both number and spatiotemporal arrangement suggests different in vivo function and recruitment of these subsets of B cells.

We have also found that HSV-2-specific antibody extracted from skin biopsies varies in concentration over the course of reactivation and healing and is present at greater concentration than antibody unrelated to HSV antigens; whereas no variability is seen in serum over the same time intervals. The variability in tissue antibody concentration corresponds to variability seen in the density of ASCs, though they do not mirror each other directly. This is congruous with observations in other conditions where a peak in ASC density is offset in time to the peak in corresponding antibody production (39, 40). Overall, the finding is congruous with a boosting of local, if not systemic, antibody secretion due to viral challenge where tissue-based antibody concentration is increased during HSV-2-reactivation and healing, and much lower in uninvolved control tissues. We strongly suspect that there is an antigen-dependent mechanism driving this process.

The role of B cells or ASCs in the inflammatory or healing process after symptomatic HSV-2 reactivation is largely unknown. Evidence from FISH and IF suggest that multiple B-cell types including mostly naïve and rarely memory cells are present. Based on the finding of B cells in multiple stages of development in biopsies and closely organized within T cell clusters, we predict that B cells are involved in antigen presentation and regulation of inflammation (41) as well as local production of antibody. Work to define the target of the antibodies produced by ASCs identified in genital skin is underway. It is possible that tissue-infiltrating memory cells may give rise to ASCs during the course of HSV-2 reactivation and healing; this needs to be tested by B-cell receptor sequencing. However, given the paucity of memory cells, it appears more likely that there is an independent mechanism by which ASCs enter inflamed tissue.

Mechanistic evaluation from human tissues is by necessity limited, and as such this study was limited in scope to those persons available and willing to undergo serial biopsies and blood draws over the course of symptomatic HSV-2 infection. In-tissue phenotyping of visualized B cells was limited by common co-expression of many canonical B cell markers. For instance, CD27 is expressed by T cells (42) which are far more numerous in the inflammatory response to HSV-2 reactivation than are B cells. Confirmation by CD27 co-staining that some of these B cells are of a memory phenotype is therefore not reliable by IF, and observation of co-expression by FISH was necessary. Likewise, CD138, or syndecan-1, is a canonical marker expressed by plasma cells and plasmablasts. It is also expressed by many non-hematopoietic-lineage cells that are highly abundant in skin including keratinocytes and fibroblasts (43). High level IgG mRNA production by FISH was therefore used in place of CD138 IF to identify potential ASCs, and the presence of CD138 co-expression was confirmed in these highly expressing IgG^+^ cells by FISH with confocal microscopy. Further investigation to identify markers of tissue infiltration and recruitment of B cells is necessary; CXCR3 and CXCL9 have been proposed (22).

In summary, we have shown the presence of B cells and ASCs present in the inflammatory infiltrate that results from HSV-2 reactivation in genital-region skin. The kinetics of tissue infiltration and egress of CD20^+^ cells, but less so IgG^+^ cells, follows that seen in T cell studies, but varies in each patient. This apparent variability may be partly due to the limited sample size available for study, however, it parallels the relative concentrations of HSV-2-specific antibody. That the concentration of HSV-2-specific antibody in tissue varies independently from the concentration of HSV-2-specific antibody in serum suggests that there are local mechanisms either leading to local production or recruitment of HSV-2-specific antibody. We propose an as of yet under-appreciated role of B cells and ASCs in the human skin-based immune response to HSV-2 reactivation.

## Methods

### Study participants

Healthy, HSV-2-seropositive adults with a history of symptomatic genital herpes were enrolled in a natural history biopsy protocol at the University of Washington Virology Research Clinic (UW-VRC) in Seattle, WA. Participants had symptoms of genital herpes for longer than 1 year and HSV-2 seropositivity was confirmed by HSV Western blot (44). Biopsies were performed by trained clinicians as described previously (5) at the time of an HSV-2 lesion, at lesion healing 10–14 days after onset and in that same location 2, 4, and 8 weeks later. A control biopsy was obtained from the upper arm or contralateral genital region skin site. Blood for peripheral blood mononuclear cells (PBMC) and plasma isolation was drawn either at enrollment or at each biopsy visit for some study participants. The University of Washington Human Subjects Review Committee reviewed and approved the protocols and all participants provided written consent.

### Materials

HSV-2 proteins gB2 and gD2 were kindly provided by Drs. G.H. Cohen and R.J. Eisenberg, gD2 was also provided by Immune Design Inc. Epstein-Barr virus (EBV) gp350 was kindly provided by Dr. A. McGuire. Stem of influenza hemagglutinin (Flu-HA) strain H1 1999 NC was expressed using vector VRC-3925 (45) in 293F cell and was kindly provided by Drs. M. Gray and L. Stamatatos.

### Immunofluorescence (IF)

Biopsy specimens were flash frozen in optimum cutting temperature (OCT) compound (Sakura Finetek) and stored at −80 °C. Slices of tissue (8 μm) perpendicular to the epidermal surface, including a cross section of the epidermis, dermal-epidermal junction (DEJ), and dermis, were prepared by cryostat sectioning and mounted on glass slides. Slides were dehydrated at room temperature for 24 hours, frozen at −80°C, and fixed in acetone prior to use. To enumerate the relative density of B cells and/or antibody in tissue biopsy sections, the following antibodies were used; mouse anti-human CD79b (1:100; Novus Biologicals), rabbit anti–human CD20 (1:100; Abcam), mouse anti-human IgG (1:100; eBioScience), sheep anti-human von-Willenbrand Factor (vWF) (FITC-conjugated, 1:5000; Abcam), and mouse anti-human CD27 (1:100; Biolegend). Prior to application of antibodies, non-specific binding was blocked with 2% bovine serum albumin, 10% casein, and 5% normal human sera, except for mouse-anti-human IgG, where human sera were omitted from the blocking buffer. TSA amplification (Invitrogen) was used for visualization of CD79b and IgG. Tissue sections were mounted in Mowiol 40-88 containing 2.5% w/v DABCO (Sigma-Aldrich). Images were captured with Nikon Eclipse Ti with NIS-Elements Software using a Hamamatsu ORCA-Flash 4.0 sCMOS camera and viewed in ImageJ/FIJI (46). CD20 positive cells were counted in ImageJ/FIJI with the cell counter plugin (47). The surface area of each biopsy was calculated in ImageJ.

### Fluorescent in situ hybridization (FISH)

To identify the presence of IgG-producing cells in biopsies and identify their spatial localization and interaction with other cell types, we performed FISH (RNAscope®2.0, ACDBio) (48). Freshly sliced, 8 μm frozen tissue sections were fixed at 4°C in 10% neutral buffered formalin, dehydrated, and treated with protease. Prepared sections were incubated with DNA-based probes for pan-IgG, CD4, CD20, CD138, PAX5, and IRF4 (ACDBio), and fluorescently tagged for visualization. After nuclear staining with DAPI (Fluka), tissues were mounted with ProLong Gold Antifade (Thermo Fisher). Images were captured with Nikon Eclipse Ti (as above) and viewed in ImageJ/FIJI. IgG RNA^+^ positive cells were counted manually over the entire slice and the surface area of each biopsy slice was calculated in ImageJ (47) to determine B-cell density. Some sections were then unmounted and stained by IF for vWF (Abcam) to determine the relative localization of B cells identified by FISH with respect to small capillaries. Colocalization of FISH probes was investigated by deconvolution (GEA/Applied Precision DeltaVision Elite) and confocal (Zeiss LSM 780 NLO) microscopy at 60x magnification, pictures were taken with a high-resolution cooled Photometrics HQ2 CCD camera.

### Preparation of biopsy extract

Five or ten freshly sliced, 10 μm biopsy cryosections were placed in a microcentrifuge tube on dry ice. To extract protein, tubes were brought to room temperature and Tissue Extraction Reagent II (Invitrogen) was added (10 μl per biopsy slice). Tube content was pipetted to mix followed by sequential vortexing (30 sec), sonication (30 sec) and pulse centrifugation. This procedure was repeated twice, then samples were flash frozen on dry ice and kept until use at –20°C. Upon thawing, samples were vortexed, sonicated and centrifuged for 5 min at 14500 x g. The supernatant was collected to measure protein, total IgG and HSV-specific antibodies via Luminex binding antibody assay.

### HSV-2 Luminex binding antibody assay

HSV-2 proteins (gB2, gD2) and control antigens (EBV gp350, Flu-HA) were coupled to MagPlex beads using an antibody coupling kit (Luminex Corp.) and stored at 4°C. To measure HSV-2 specific antibody, MagPlex beads were incubated with blocking buffer (5% Blotting-Grade Blocker (Bio-Rad), 0.05% Tween-20 (Sigma), phosphate buffered saline (PBS)) to minimize non-specific binding. Beads were then washed and mixed with serial dilutions of biopsy extracts or serum samples in assay buffer (Pierce™ Protein-Free (PBS) Blocking Buffer, 1% Blotting-Grade Blocker, and 0.05% Tween-20). Sera pooled from ten HSV-2 seropositive and ten HSV-2 seronegative donors were used as positive and negative controls, respectively. After incubation with biopsy extracts or serum samples, MagPlex beads were washed with PBST (PBS, 0.05% Tween-20) and incubated with anti-human IgG Fc-PE (Southern Biotech). Finally, beads were washed 3 times and resuspended in PBS containing 1% BSA and 0.05% Tween-20. Median fluorescence intensity (MFI) by PE fluorescence for each bead type was collected on Luminex 200 instruments (Luminex Corp.) operated by MagPlex software (Hitachi) or xPonent Software (Luminex Corp.). The level of background was assigned by the MFI of antigen-conjugated beads incubated first with buffer (in place of serum), then with secondary antibody. Background MFI values for each antigen were subtracted from experimental measurements. The MFI of the biopsy extracts was normalized to the surface area of each biopsy slice (as measured in adjacent slices) to account for variation in size and protein composition of biopsy specimens. Area under the curve (AUC) for serum titration experiments was calculated in GraphPad Prism.

### HSV-2 quantitative PCR

DNA from biopsies and genital swabs was extracted using tissue kits (EZ1; QIAGEN) from 60–80 μm cross-sectioned samples from each biopsy specimen as previously described (49, 50), including subtype-specific typing by real-time PCR (51). HSV-2 DNA copy numbers were normalized to 1×10^6^ cells using β-globin copy numbers. Primers used were 5ʹ-TGAAGGCTCATGGCAAGAAA-3ʹ and 5ʹ-GCTCACTCAGTGTGGCAAAGG-3ʹ with 5ʹ-TCCAGGTGAGCCAGGCCATCACTA-3ʹ as a probe. Detection limits of HSV-2 DNA quantification were one copy per 50,000 cells in tissue, as previously published (34, 49).

### Statistics

All statistics and data analysis were performed using R Studio (R 3.4.1, R Core Team, Vienna, Austria) or GraphPad Prism version 8.2.0 for Mac (GraphPad Software, California, USA). To compare cell density and antibody concentration between biopsy time points the Friedman test with Dunn’s correction for multiple comparisons was used. To test the observation of greatest B cell and ASC density the maximum of lesion or newly healed was compared to the maximum at 2, 4 or 8 weeks after healing by a paired Wilcoxon test. Correlation between the density of cell types was tested by non-parametric Spearman correlation due to the presence of multiple biopsies without B cells. Comparison of serum antibody titer over time was made by area under the curve (AUC) with the Friedman test with Dunn’s correction for multiple comparisons. Statistical significance was defined as adjusted, two-sided p value ≤0.05.

### Study approval

The natural history biopsy protocol was approved by the University of Washington Institutional Review Board (IRB) and written, informed consent was obtained from each participant.

## Author contributions

Study design: ESF, AMS, JZ, LC

Tissue biopsy analysis: ESF, AMS, AK, KP, RMB, SSC

Participant recruitment and clinical guidance: CJ, AW

Data analysis: ESF, AMS, RMB, JS, RR, PTE, JZ, TP

Manuscript preparation: ESF, AMS, JZ, LC

All authors reviewed and approved of the manuscript.

## Acknowledgements

The authors would like to express their appreciation to Mindy Miner for her assistance in editing and formatting the final manuscript and to David McDonald for assistance in confocal and deconvolution microscopy.

---

**Supplementary Table 1.** Participant demographics and HSV clinical history.

| Participant | Age | Gender | Location of HSV lesion | Serologic status | Site of control biopsy |
| --- | --- | --- | --- | --- | --- |
| 1 | 47 | F | Perineal | HSV-1&2 | Contra |
| 2 | 63 | F | Buttock | HSV-1&2 | Contra |
| 3 | 61 | F | Buttock | HSV-2 | Arm |
| 4 | 46 | F | Labia | HSV-2 | Arm |
| 5 | 26 | M | Mons | HSV-1&2 | Arm |
| 6 | 32 | F | Labia | HSV-2 | Contra |
| 7 | 57 | F | Buttock | HSV-1&2 | Contra |
| 8 | 51 | F | Buttock | HSV-2 | Contra |
| 9 | 49 | F | Buttock | HSV-2 | Arm |
| 10 | 47 | F | Buttock | HSV-1&2 | Contra |
| 11 | 59 | F | Buttock | HSV-2 | Arm |
| 12 | 58 | F | Perianal | HSV-2 | Arm |
| 13 | 53 | M | Buttock | HSV-2 | Contra |
| 14 | 18 | F | Mons | HSV-2 | Contra |
| 15 | 66 | F | Buttock | HSV-1&2 | Arm |
| 16 | 33 | F | Buttock | HSV-2 | Contra |

**Supplementary Figure 1.**
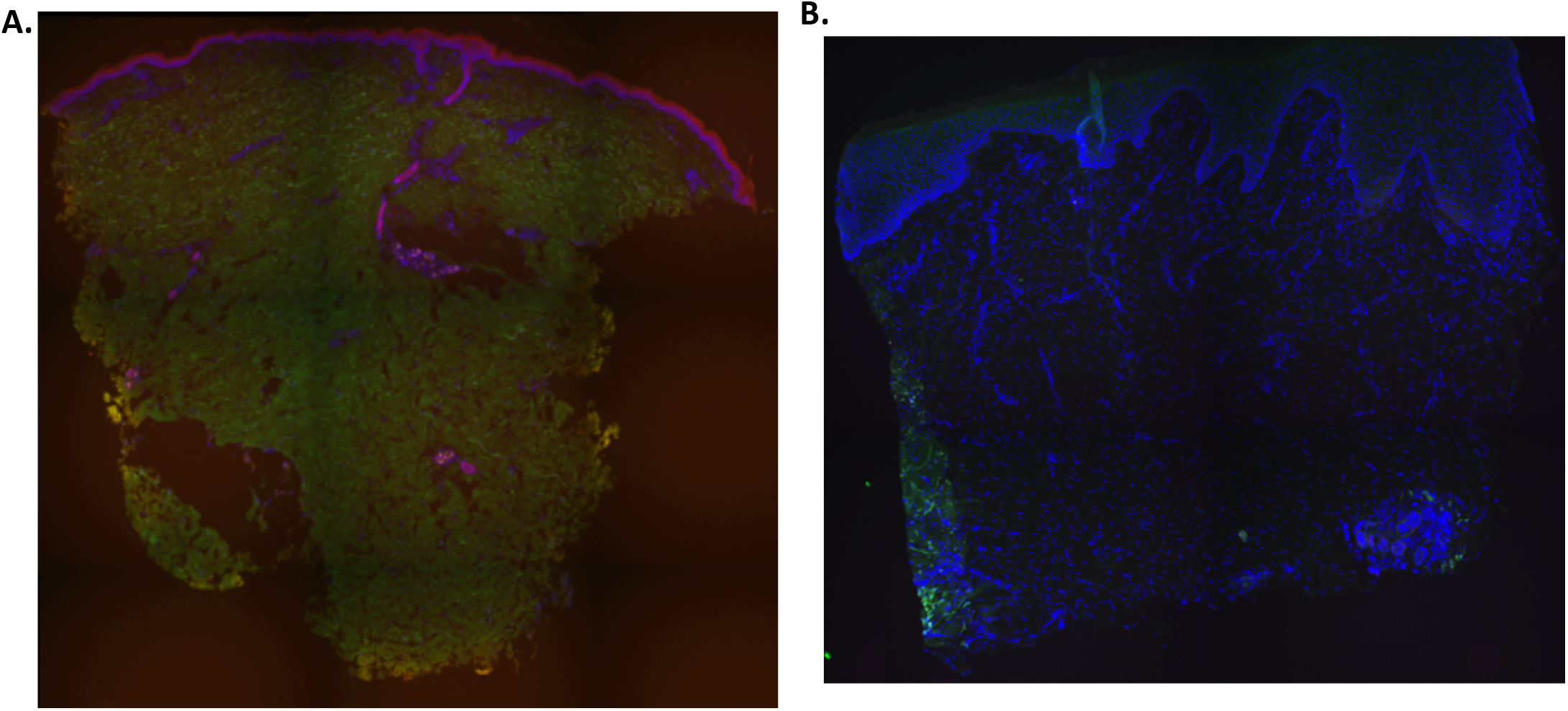
Isotype control (A) and isolated primary antibody (B), (no secondary antibody) for IgG IFA.

**Supplementary Table 2.** Area under the curve (AUC) of serum samples collected concurrently with genital skin biopsies.

| Participant 1 | Lesion/<br>Control | Healing | 2 wph | 4 wph | 8 wph |
| --- | --- | --- | --- | --- | --- |
| gB2 | 32797 | 34319 | 33089 | 33492 | 33103 |
| gD2 | 23328 | 25318 | 23833 | 23665 | 23532 |
| Flu-HA | 9890 | 10731 | 9931 | 9890 | 9321 |
| EBV gp350 | 654 | 744 | 657 | 637 | 582 |
| Participant 6 | Lesion/<br>Control | Healing | 2 wph | 4 wph | 8 wph |
| gB2 | 26953 | 26821 | 24505 | 26847 | 26046 |
| gD2 | 25590 | 25790 | 23541 | 25383 | 24838 |
| Flu-HA | 18527 | 18100 | 16195 | 17926 | 17512 |
| EBV gp350 | 19714 | 19340 | 17472 | 19159 | 19110 |

**Supplementary Figure 2.**
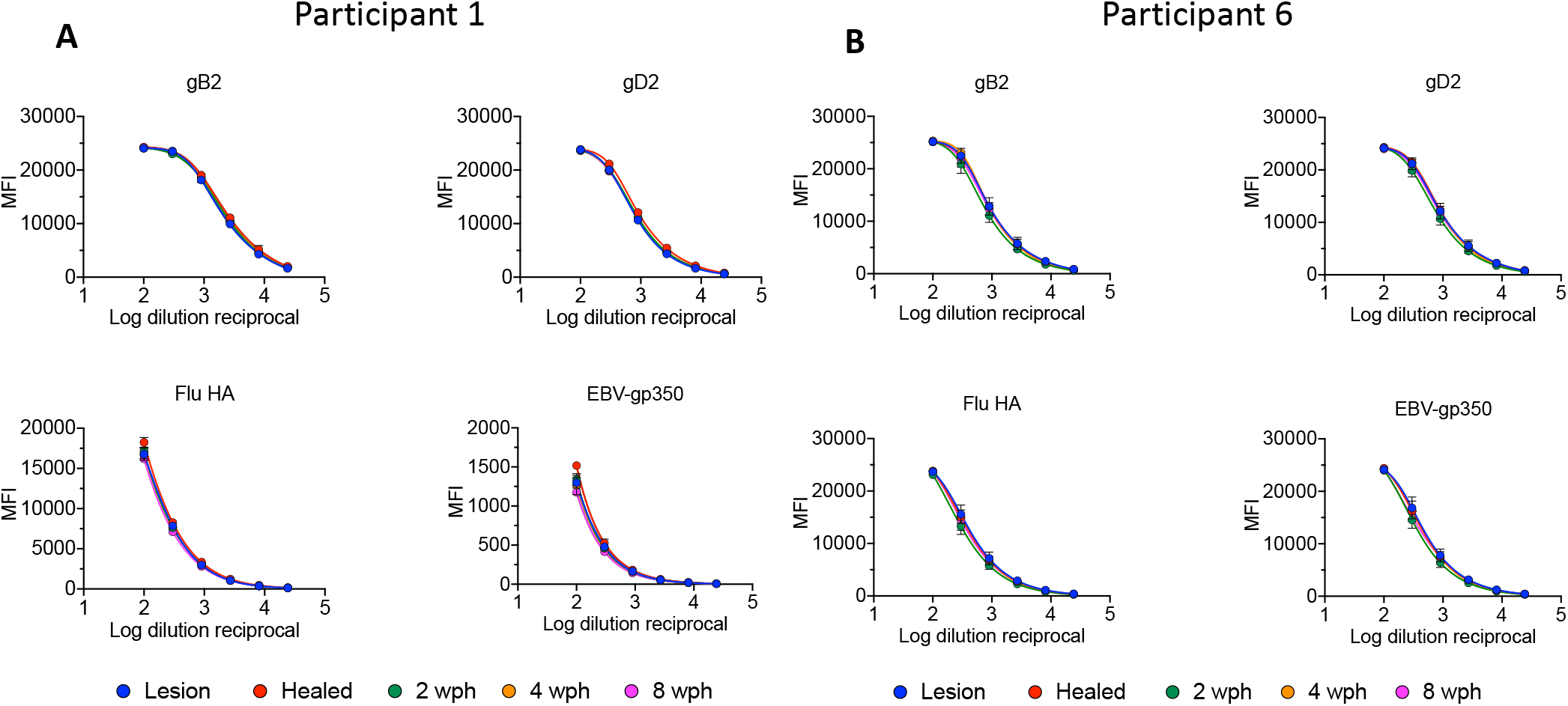
Serum levels of HSV-2 specific antibody do not change during HSV-2 reactivation. IgG levels are expressed as units of Median Fluorescence Intensity (MFI) over serum dilution. Serum samples were collected concurrently with genital skin biopsies at: lesion (blue), healing (red), 2 wph (green), 4 wph (orange) and 8 wph (magenta) time points. (**A**) Participant 1; (**B**) Participant 6.

**Supplementary Figure 3.**
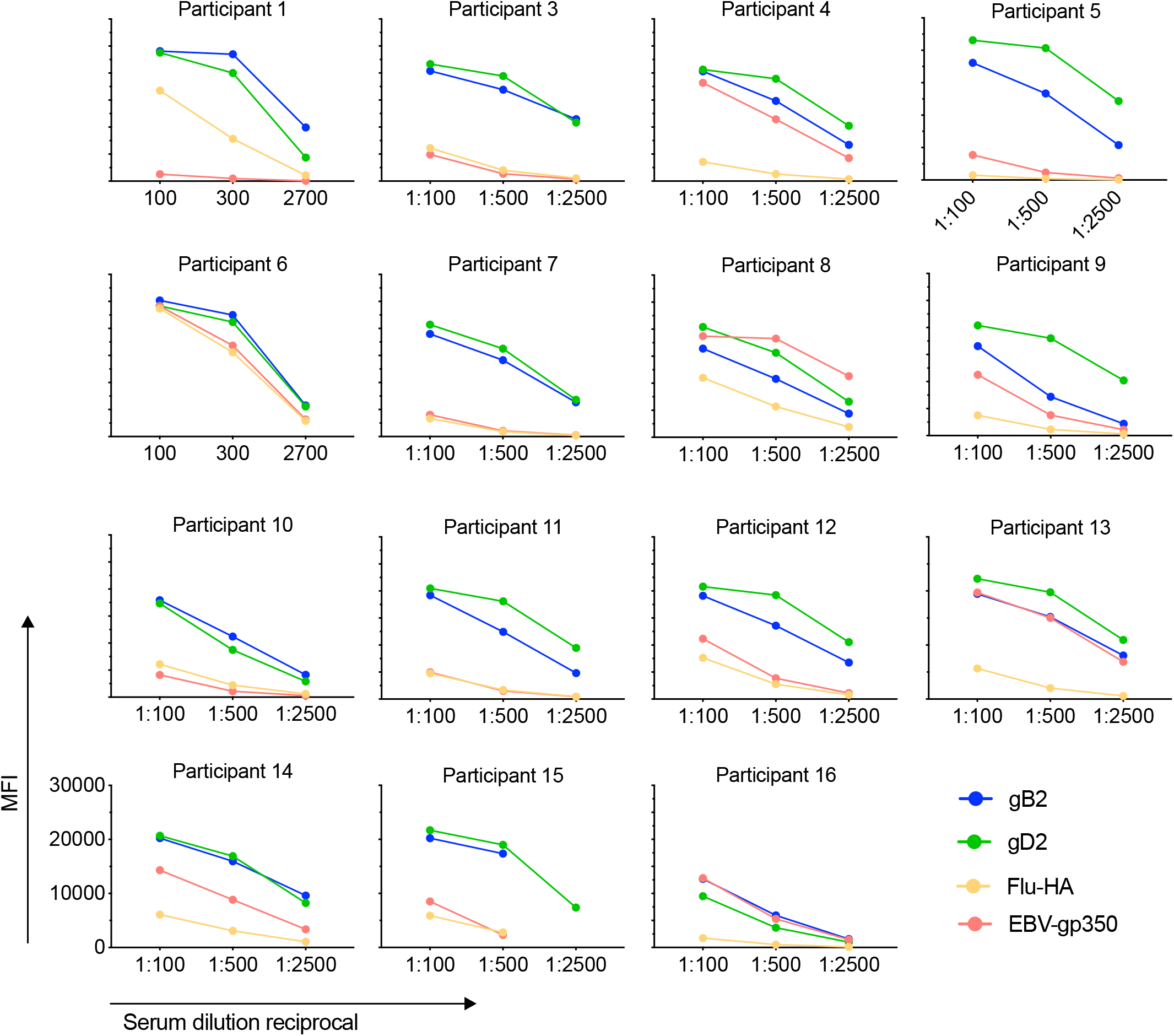
HSV-2, EBV-gp350 and Flu-HA specific IgG in serum samples collected at participant enrollment.

**Supplementary Table 3.**
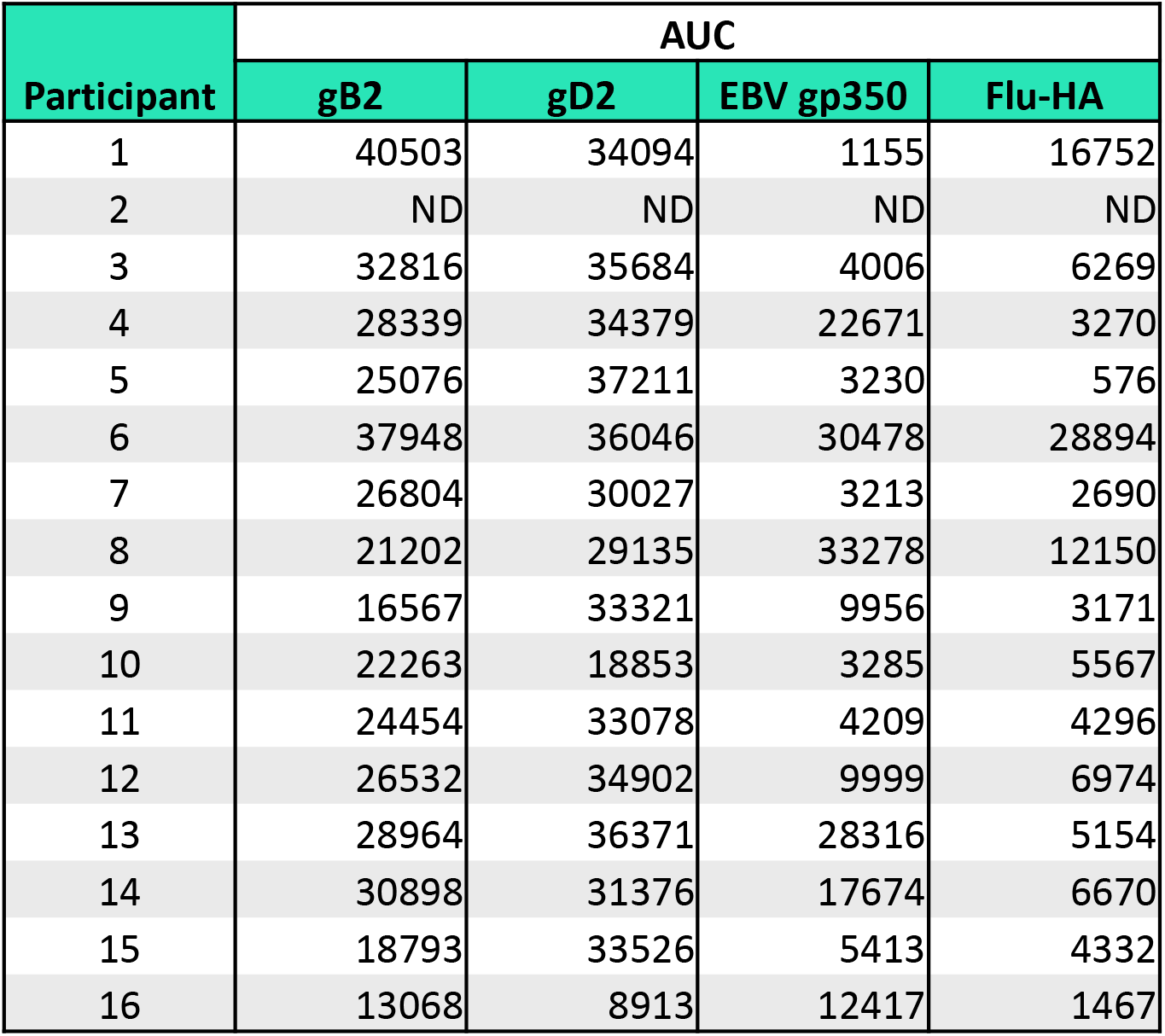
Area under the curve (AUC) measurements of serum samples collected at participant enrollment.

| Participant | AUC |  |  |  |
| --- | --- | --- | --- | --- |
|  | gB2 | gD2 | EBV gp350 | Flu-HA |
| 1 | 40503 | 34094 | 1155 | 16752 |
| 2 | ND | ND | ND | ND |
| 3 | 32816 | 35684 | 4006 | 6269 |
| 4 | 28339 | 34379 | 22671 | 3270 |
| 5 | 25076 | 37211 | 3230 | 576 |
| 6 | 37948 | 36046 | 30478 | 28894 |
| 7 | 26804 | 30027 | 3213 | 2690 |
| 8 | 21202 | 29135 | 33278 | 12150 |
| 9 | 16567 | 33321 | 9956 | 3171 |
| 10 | 22263 | 18853 | 3285 | 5567 |
| 11 | 24454 | 33078 | 4209 | 4296 |
| 12 | 26532 | 34902 | 9999 | 6974 |
| 13 | 28964 | 36371 | 28316 | 5154 |
| 14 | 30898 | 31376 | 17674 | 6670 |
| 15 | 18793 | 33526 | 5413 | 4332 |
| 16 | 13068 | 8913 | 12417 | 1467 |

## References

1. Wald A, Zeh J, Selke S, Ashley RL, Corey L. Virologic characteristics of subclinical and symptomatic genital herpes infections. N. Engl. J. Med. 1995;333(12):770–775.

2. Benedetti J, Corey L, Ashley R. Recurrence rates in genital herpes after symptomatic first-episode infection. Ann. Intern. Med. 1994;121(11):847–854.

3. Schiffer JT et al. Mathematical modeling predicts that increased HSV-2 shedding in HIV-1 infected persons is due to poor immunologic control in ganglia and genital mucosa. PLoS One 2016;11(6):e0155124.

4. Schiffer JT et al. Mucosal host immune response predicts the severity and duration of herpes simplex virus-2 genital tract shedding episodes.. Proc. Natl. Acad. Sci. U. S. A. 2010;107(44):18973–18978.

5. Zhu J et al. Virus-specific CD8+ T cells accumulate near sensory nerve endings in genital skin during subclinical HSV-2 reactivation. J. Exp. Med. 2007;204(3):595–603.

6. Iijima N et al. Dendritic cells and B cells maximize mucosal Th1 memory response to herpes simplex virus.. J. Exp. Med. 2008;205(13):3041–3052.

7. Zhu J et al. Persistence of HIV-1 receptor-positive cells after HSV-2 reactivation is a potential mechanism for increased HIV-1 acquisition.. Nat. Med. 2009;15(8):886–892.

8. Zhu J et al. Immune surveillance by CD8αα+ skin-resident T cells in human herpes virus infection.. Nature 2013;497(7450):494–7.

9. Schiffer JT et al. Rapid localized spread and immunologic containment define herpes simplex virus-2 reactivation in the human genital tract. Elife 2013;2:1–28.

10. Iijima N, Iwasaki A. A local macrophage chemokine network sustains protective tissue-resident memory CD4 T cells. Science 2014;346(6205):93–98.

11. Mertz GJ et al. Frequency of acquisition of first-episode genital infection with herpes simplex virus from symptomatic and asymptomatic source contacts.. Sex. Transm. Dis. 1985;12(1):33–9.

12. Cairns TM et al. Dissection of the antibody response against herpes simplex virus glycoproteins in naturally infected humans. J. Virol. 2014;88(21):12612–12622.

13. Flechtner JB et al. Immune responses elicited by the GEN-003 candidate HSV-2 therapeutic vaccine in a randomized controlled dose-ranging phase 1/2a trial. Vaccine 2016;34(44):5314–5320.

14. Bernstein DI et al. Therapeutic vaccine for genital herpes simplex virus-2 infection: Findings from a randomized trial. J. Infect. Dis. 2017;215(6):856–864.

15. Wald A et al. Therapeutic HSV-2 vaccine (GEN003) results in durable reduction in genital lesions at 1 year. Open Forum Infect. Dis. 2014;1(Suppl 1):S55–S56.

16. Gilbert PB et al. Antibody to HSV gD peptide induced by vaccination does not protect against HSV-2 infection in HSV-2 seronegative women. PLoS One 2017;12(5):1–15.

17. Burn C et al. An HSV-2 single-cycle candidate vaccine deleted in glycoprotein D, ΔgD-2, protects male mice from lethal skin challenge with clinical isolates of HSV-1 and HSV-2. J. Infect. Dis. 2017;(January):1–5.

18. Jiang Y et al. Maternal antiviral immunoglobulin accumulates in neural tissue of neonates to prevent HSV neurological disease. MBio 2017;8(4):1–14.

19. Criscuolo E et al. Cell-to-cell spread-blocking activity is extremely limited in the sera of HSV-1 and HSV-2 infected subjects.. J. Virol. 2019;JVI.00070-19.

20. Harandi AM, Svennerholm B, Holmgren J, Eriksson K. Differential roles of B cells and IFN-??-secreting CD4+ T cells in innate and adaptive immune control of genital herpes simplex virus type 2 infection in mice. J. Gen. Virol. 2001;82(4):845–853.

21. Schenkel JM et al. Resident memory CD8 T cells trigger protective innate and adaptive immune responses. Science 2014;346(6205):98–101.

22. Oh JE et al. Migrant memory B cells secrete luminal antibody in the vagina. Nature 2019;571:122–126.

23. Wilson RP et al. IgM plasma cells reside in healthy skin and accumulate with chronic inflammation. J. Invest. Dermatol. 2019;139(12):2477–2487.

24. Mandal A et al. Cell and fluid sampling microneedle patches for monitoring skin-resident immunity. Sci. Transl. Med. 2018;10(468). doi:10.1126/scitranslmed.aar2227

25. Tabib T, Morse C, Wang T, Chen W, Lafyatis R. SFRP2/DPP4 and FMO1/LSP1 define major fibroblast populations in human skin. J. Invest. Dermatol. 2018;138(4):802–810.

26. Lemos MP et al. In men at risk of HIV infection, IgM, IgG1, IgG3, and IgA reach the human foreskin epidermis. Mucosal Immunol. 2015;(April):1–11.

27. Cunningham AL, Turner RR, Miller AC, Para MF, Merigan TC. Evolution of recurrent herpes simplex lesions: an immunohistologic study. J. Clin. Invest. 1985;75(1):226–233.

28. Egbuniwe IU, Karagiannis SN, Nestle FO, Lacy KE. Revisiting the role of B cells in skin immune surveillance. Trends Immunol. 2015;36(2):102–111.

29. Geherin S a et al. The skin, a novel niche for recirculating B cells. J. Immunol. 2012;188(12):6027–6035.

30. Ginaldi L, De Martinis M, D’Ostilio A, Marini L, Quaglino D. Changes in antigen expression on B lymphocytes during HIV infection. Pathobiology 1998;66(1):17–23.

31. Stevens RH, Askonas BA, Welstead JL. Immunoglobulin heavy chain mRNA in mitogen-stimulate B cells. Eur. J. Immunol. 1975;5(1):47–53.

32. Coronella J a, Telleman P, Truong TD, Ylera F, Junghans RP. Amplification of IgG VH and VL (Fab) from single human plasma cells and B cells.. Nucleic Acids Res. 2000;28(20):E85.

33. Ellebedy AH et al. Defining antigen-specific plasmablast and memory B cell subsets in human blood after viral infection or vaccination. Nat. Immunol. 2016;17(10):1226–1234.

34. Peng T et al. An effector phenotype of CD8+ T cells at the junction epithelium during clinical quiescence of herpes simplex virus 2 infection. J. Virol. 2012;86(19):10587–10596.

35. Douglas RG, Couch RB. Infection and Recurrent Herpes Labialis in Humans 11970;(2):289–295.

36. Friedman MG, Kimmel N. Herpes simplex virus-specific serum immunoglobulin A: Detection in patients with primary or recurrent herpes infections and in healthy adults. Infect. Immun. 1982;37(1):374–377.

37. Juto P, Settergren B. Specific serum IgA, IgG and IgM antibody determination by a modified indirect ELISA-technique in primary and recurrent herpes simplex virus infection. J. Virol. Methods 1988;20(1):45–55.

38. Woodman CB et al. The relative infrequency and low levels of neutralising and immunoprecipitating antibody to herpes simplex viruses types 1 and 2 in patients with a history of recurrent herpes genitalis.. Med. Microbiol. Immunol. 1983;171(4):243–250.

39. Blanchard-Rohner G, Pulickal AS, Jol-van Der Zijde CM, Snape MD, Pollard AJ. Appearance of peripheral blood plasma cells and memory B cells in a primary and secondary immune response in humans. Blood 2009;114(24):4998–5002.

40. He XS et al. Plasmablast-derived polyclonal antibody response after influenza vaccination. J. Immunol. Methods 2011;365(1–2):67–75.

41. Griss J et al. 479 B cells sustain inflammation and predict response to immune checkpoint blockade in human melanoma. J. Invest. Dermatol. 2019;139(9):S297.

42. Hintzen RQ et al. Regulation of CD27 expression on subsets of mature T-lymphocytes.. J. Immunol. 1993;151(5):2426–35.

43. O’Connell FP, Pinkus JL, Pinkus GS. CD138 (Syndecan-1), a plasma cell marker immunohistochemical profile in hematopoietic and nonhematopoietic neoplasms. Am. J. Clin. Pathol. 2004;121(2):254–263.

44. Ashley RL, Militoni J, Lee F, Nahmias A, Corey L. Comparison of Western blot (immunoblot) and glycoprotein G-specific immunodot enzyme assay for detecting antibodies to herpes simplex virus types 1 and 2 in human sera. J. Clin. Microbiol. 1988;26(4):662–667.

45. Lingwood D et al. Structural and genetic basis for development of broadly neutralizing influenza antibodies. Nature 2012;489(7417):566–570.

46. Schindelin J et al. Fiji: An open-source platform for biological-image analysis. Nat. Methods 2012;9(7):676–682.

47. Diem K et al. Image analysis for accurately counting CD4+ and CD8+ T cells in human tissue. J. Virol. Methods 2015;222:117–121.

48. Wang F et al. RNAscope: A novel in situ RNA analysis platform for formalin-fixed, paraffin-embedded tissues. J. Mol. Diagnostics 2012;14(1):22–29.

49. Magaret AS, Wald A, Huang M-L, Selke S, Corey L. Optimizing PCR positivity criterion for detection of herpes simplex virus DNA on skin and mucosa. J. Clin. Microbiol. 2007;45(5):1618–1620.

50. Wald A, Huang M-L, Carrell D, Selke S, Corey L. Polymerase chain reaction for detection of herpes simplex virus (HSV) DNA on mucosal surfaces: comparison with HSV isolation in cell culture.. J. Infect. Dis. 2003;188(9):1345–51.

51. Corey L, Huang M-L, Selke S, Wald A. Differentiation of herpes simplex virus types 1 and 2 in clinical samples by a real-time taqman PCR assay. J. Med. Virol. 2005;76:350–355.

